# HNF4α is a target of the Wnt/β-catenin pathway and regulates colorectal carcinogenesis

**DOI:** 10.1101/2024.03.25.586542

**Authors:** Lei Sang, Weiyu Bai, Chenglu Lu, Rui Dong, Qinggang Hao, Yingru Zhang, Rongyuan Sun, Junling Shen, Yan Sun, Jianwei Sun

**Author notes:** These authors contributed equally to this work (Lei Sang, Weiyu Bai, Chenglu Lu, Rui Dong). Correspondence to Jianwei Sun or Yan Sun or Junling Shen.

## Abstract

Hepatocyte nuclear factor 4alpha (HNF4α) is a transcription factor involved in liver function. Dysregulation of HNF4α leads to hepatocarcinogenesis. However, the role and mechanism of HNF4α in colorectal cancer is still unknown. Here we demonstrate that HNF4α is upregulated in colorectal cancers. and HNF4α upregulation promotes colorectal tumorigenesis. Notably, expression levels of HNF4α are positively correlated with the Wnt/β-catenin signaling pathway in colorectal cancer patients. Further, we showed that HNF4α is transcriptionally activated by Wnt/β-catenin/TCF3 and is at least partially responsible for the oncognesis activity of the Wnt signaling in colorectal cancer. Our findings indicate that HNF4α is a direct target of that Wnt/β-catenin pathway and could play an important role in colorectal carcinogenesis.

**Author Summary:** Here we demonstrate that HNF4α is upregulated in colorectal cancers and HNF4α upregulation promotes colorectal tumorigenesis. Notably, expression level of HNF4α are positively correlated with the Wnt/β-catenin pathway in colorectal cancer patients. Further, we showed that HNF4α is transcriptionally activated by Wnt/β-catenin/TCF3 and is at least partially responsible for the oncognesis activity of the Wnt signaling pathway in colorectal cancer.

## Introduction

Colorectal cancer (CRC) is graded as one of the most common cancers and accounts for the second leading cause of cancer deaths worldwide. Despite an overall decreasing incidence of the disease, early-onset colorectal cancer is a growing concern and rapidly rising in those under the age of 50 over the last two decades[1–3]. While a number of major targets have been identified during the course of CRC development such as Ras[4], Tp53[5], and APC/β-catenin[6], the pathogenesis of CRC still remains elusive. Identification of new molecules that are involved in CRC could lead to early detection, prevention and therapeutic intervention.

Hepatocyte Nuclear Factor 4 Alpha (HNF4α), a master transcription factor, is predominantly expressed in the liver, kidney, intestine and endocrine pancreas, and is necessary for liver development and function[7, 8]. However, accumulating evidence shows its inhibitory role in the liver cancer. The HNF4α mutations at G79C, F83C and M125I (Zn-finger DNA-binding domain region) were detected in liver cancer and were believed to trigger liver tumorigenesis[9]. Previous studies revealed the contradictory role of HNF4α in colorectal carcinogenesis[10]. The regulation of HNF4α in CRC is largely unknown. Further investigation of the role and the specific regulation mechanism of HNF4α in the development and progression of CRC is of great significance for establishing HNF4α as a diagnostic/prognostic marker and therapeutic target in CRC.

The Wnt/β-catenin signaling pathway is a recognized driver of colorectal carcinogenesis and the defect of this cascade occurs in 70–80% of CRC [11, 12]. Activation of the Wnt pathway induces β-catenin translocation into the nucleus where it functions as a transcriptional coactivator of TCF /LEF family factors[13]. While several molecules that regulate this cascade have been identified including APC, GSK3β and AXIN, the role and regulation of Wnt/β-catenin activation in CRC is still not completely understood.

Here, we demonstrated that HNF4α was upregulated in CRC and promoted colorectal tumorigenesis. HNF4α levels were positively correlated with and directly regulated by the Wnt/β-catenin-TCF3 signaling pathway. We further showed that HNF4α is a positive regulator of the Wnt/β-catenin and that HNF4α overexpression promotes β-catenin nuclear localization. Our results suggested that HNF4α is a potential prognostic marker for the Wnt/β-catenin-related CRC and that HNF4α forms a feedback loop with Wnt/β-catenin to control colorectal tumorigenesis.

## Results

### 1. HNF4 α promotes colorectal carcinogenesis

Recently, we found that the loss of *nhr-14* gene in *c. elegans* resulted in the dysregulation of DNA damage-induced response [14], which usually leads to cancer development [15–18]. Thus, we examined HNF4α expression levels in different cancers and matched paired normal tissues. TCGA database analysis revealed upregulation of HNF4α in Colon adenocarcinoma (COAD), Rectum adenocarcinoma (READ) and Pancreatic adenocarcinoma (PAAD), suggesting that HNF4α may play an important role in gastrointestinal tumorigenesis, especially in colorectal carcinogenesis (Figure 1A). We further analyzed the HNF4α protein levels in 59 paired colon cancer and normal tissues and found significant increase of HNF4α in 43 tumors (Figure 1B and 1C). Immunohistochemical staining of CRC microarray also revealed remarkable upregulation of HNF4α in cancer tissues (n=244) when compared to normal colon tissues (n=99) (Figure 1D). These results indicate that overexpression of HNF4α is a recurrent event in CRC and could play a significant role in this malignancy.

**Figure 1.**
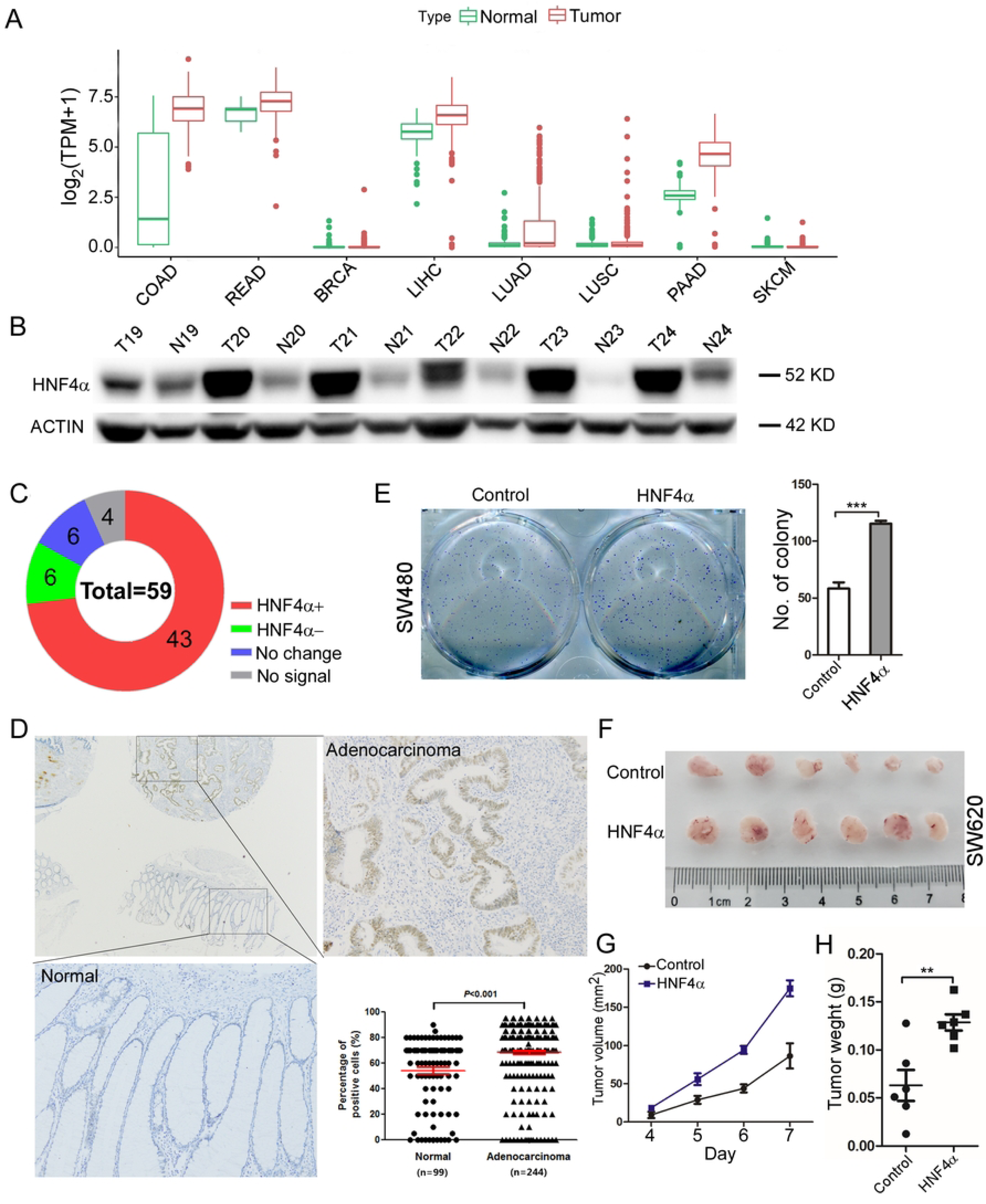
HNF4α is frequently upregulated in CRC and plays an oncogenic role during colorectal tumorigenesis. A. TCGA database analysis of HNF4α expression in different tumor types. B. Western blot analysis of HNF4α protein level in CRC tumor (T) and corresponding adjacent normal (N) tissues. C. Schematic diagram of HNF4α expression in 59 pairs of CRC tumor and normal tissues. D. Immunohistochemical staining of CRC tissue microarray (TMA) with an antibody against HNF4α and statistical analysis of HNF4α expression levels in the CRC TMA. E. Effects of ectopic expression of HNF4α on colony formation in SW480 cells. F. Tumor volume of mouse xenografts established with parental (6 mice) and HNF4α overexpressing (6 mice) SW480. G. Growth curve reflects the effect HNF4α overexpression on tumor growth in xenograft mouse model. H. Effects HNF4α overexpression on tumor weight. * P<0.05, ** P<0.01 and *** P<0.001.

Previous studies have shown that HNF4α could behave as either a tumor suppressor or an oncogene largely depending on the tumor type and organ[10, 18, 19]. Thus, we next investigated the function of HNF4α in CRC by stable transfection of HNF4α into SW480 and SW620 colon cancer cell lines. We found that ectopic expression of HNF4α significantly increased colony formation (Figure 1E). Xenograft mouse experiments also showed that overexpression of HNF4α considerably promoted the tumor growth and increased the tumor volume and weight (Figure F-H). These findings suggest that HNF4α acts as an oncogene and could be a driver in CRC.

### 2. HNF4α is positively correlated with and is regulated by the Wnt/β-catenin signaling pathway

To examine the mechanism by which HNF4α is upregulated in CRC, we downloaded the COAD and READ data from TCGA database and performed GESA analysis. Among the enriched HNF4α-correlated signaling pathway, oncogenic Wnt/β-catenin and MYC pathways were significantly and positively associated with HNF4α expression (Figure 2A). It has been reported that the WNT/β-catenin transcriptionally induces c-Myc expression to promote colorectal tumorigenesis [20]. Thus, we primarily focused on the relationship between HNF4α and the Wnt/β-catenin pathway. We also analyzed the correlation between β-catenin and HNF4α, and the result indicated that β-catenin positively associated with HNF4α expression levels in TCGA database (Figure 2B). Then we transfected SW480 cells with shRNA-HNF4α and control shRNA and found no expression change of CTNB1 at mRNA or protein levels between HNF4α-knockdown and control cells (data not shown). Conversely, SW480 cells after transfection with shRNA-CTNB1 expressed significant low levels of HNF4α mRNA when compared to the cells treated with control shRNA (Figure 2C). Moreover, we transfected shRNA-CTNB1 into other 2 (SW620 and DLD1) colon cancer cell lines. Western blot analysis showed that knockdown of β-catenin significantly reduced the protein level of HNF4α in SW480, SW620 and DLD1 cell lines (Figure 2D). These results suggest that HNF4α is regulated by the WNT /β-catenin pathway at the transcription level.

**Figure 2.**
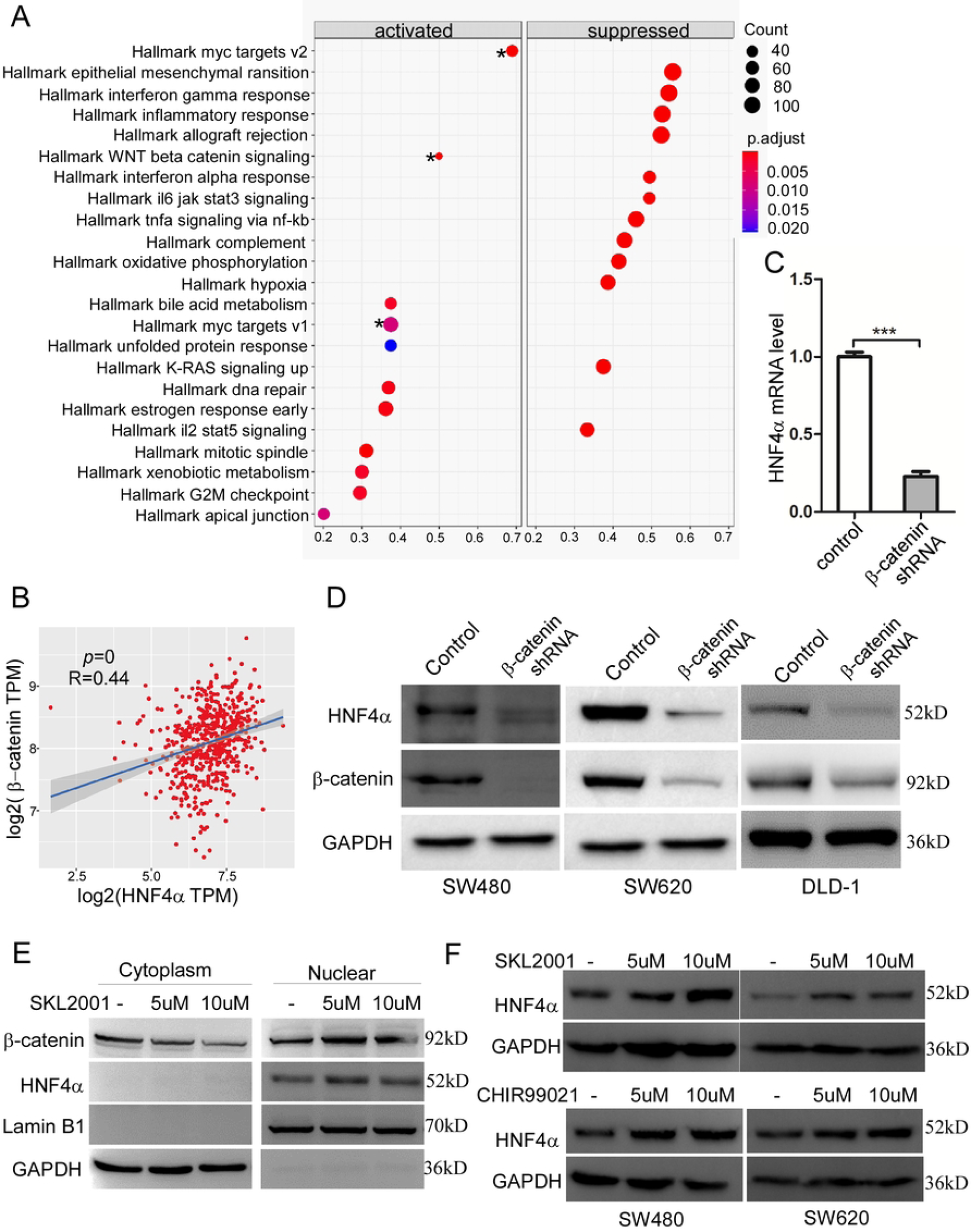
HNF4α is a target of Wnt/β-catenin. A. GESA analysis of HNF4α in TCGA database, * indicates the significant correlation of HNF4α with Wnt and Myc pathways. B. Correlations analysis between HNF4α and β-catenin mRNA level in TCGA database. C. Q-PCR analysis of the effect of β-catenin knockdown on HNF4α mRNA level. D. Western blotting analysis of the effect of β-catenin knockdown on HNF4α protein expression in 3 different colon cancer cell lines. E. Western blotting analysis of the effect of SKL2001 on subcellular location and expression of β-catenin and HNF4α. F. Western blotting analysis of the effect of SKL2001 and Chir-99021, 2 activators of WNT/β-catenin signaling, on HNF4α expression in SW480 and SW620 cell lines. ***p<0.001.

To further support this conclusion, we treated colon cancer cells with SKL2001, an activator of the Wnt/β-catenin pathway by disrupting Axin/β-catenin interaction, at different doses for 12h. Western blot analysis of cellular fractionation revealed significant nuclear accumulation of β-catenin and increased expression of HNF4α following SKL2001 treatment (Figure 2E-F). In addition, we treated SW480 and SW620 cells with Chir-99021, another β-catenin activator by selectively inhibiting GSK-3α/β, and found upregulation of HNF4α in both cell lines after Chir-99021treatment (Figure 2F). These results further confirmed that β-catenin is an activator of HNF4α.

### 3. Ectopic expression of HNF4α rescues β-catenin knockdown in tumor growth

As HNF4α is a downstream target of the Wnt//β-catenin pathway, we next invested whether HNF4α mediates the role of the Wnt/β-catenin axis in colon carcinogenesis. SW620 cells were stably transfected with β-catenin shRNA or β-catenin shRNA and HNF4α overexpression, respectively. The cells treated with vector alone were used as control (Figure 3A). These cells were subcutaneously injected to nude mice. As expected, knockdown of β-catenin inhibited tumor growth and reduced tumor volume and weight (Figure 3B-E). Notably, ectopic expression of HNF4α largely overrode these phenotypes caused by β-catenin knockdown (Figure 3B-E). These findings further indicate that HNF4α is a major target of β-catenin in colon cancer.

**Figure 3.**
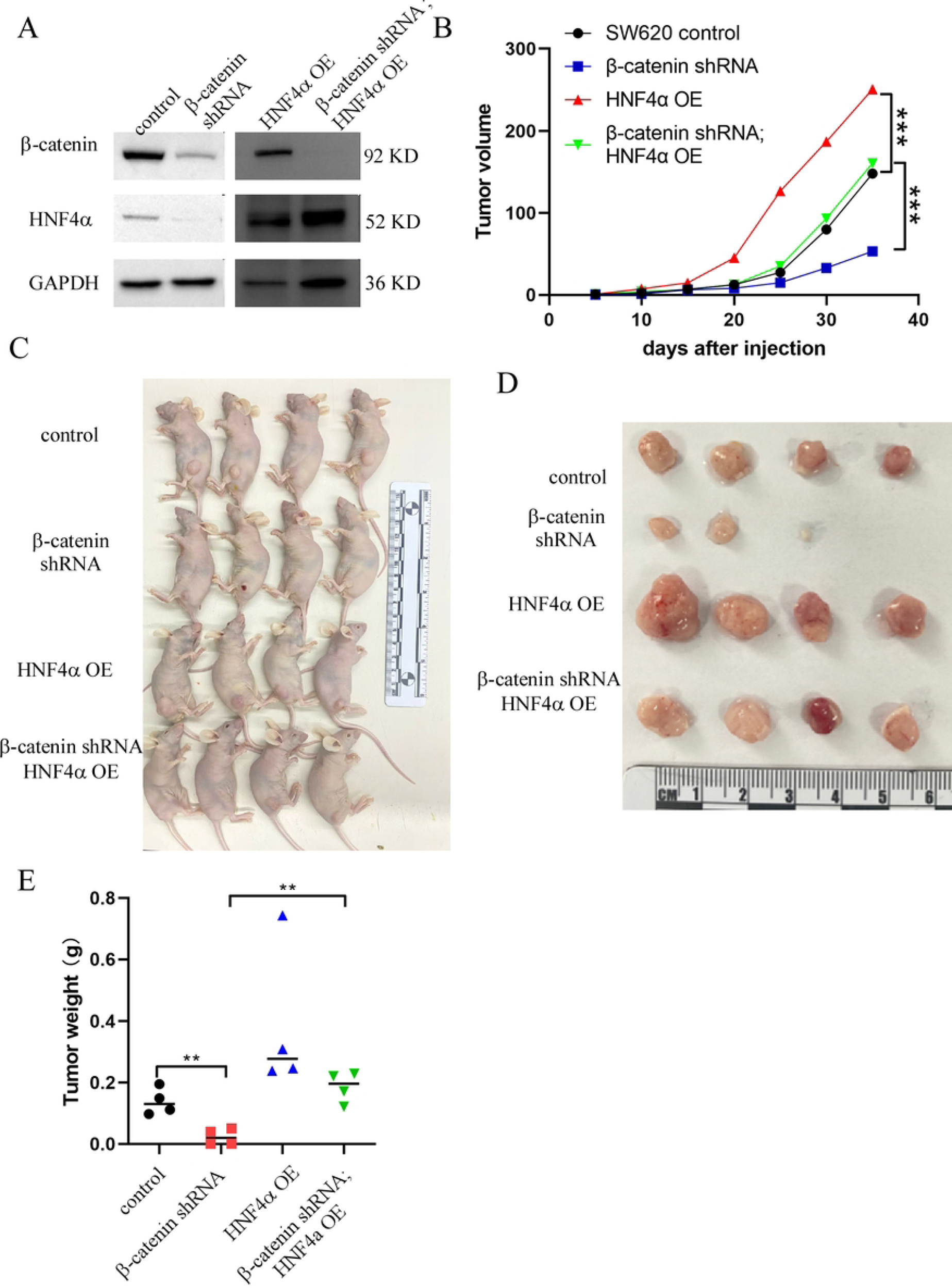
HNF4α mediates Wnt/β-catenin function in colon tumorigenesis. A. Western blotting analysis of β-catenin and HNF4α expression in SW620 cells transfected with β-catenin shRNA, β-catenin shRNA/HNF4α. B. Effect of β-catenin knockdown, β-catenin shRNA/HNF4**α** on tumor growth. C. Representative images of the effects of β-catenin shRNA, β-catenin shRNA/HNF4**α** on tumor growth. D. Effect of β-catenin shRNA, β-catenin shRNA/HNF4**α** on tumor size. E. Effect of β-catenin shRNA, β-catenin shRNA/HNF4**α** on tumor weight. * p <0 .05, ** p<0.01, ***p<0.001.

### 4. TCF3 (TCF7L1) promotes HNF4α expression

β-catenin is a transcription co-activator of TCF/LEF transcription factor (TF) family. To further understand the molecular mechanisms by which wnt/ β -catenin induces HNF4α transcription, we first analyzed the expression relationship between HNF4α and TCF7(TCF-1), LEF1, TCF7L1(TCF-3), TCF7l2(TCF-4), which are TCF/LEF family members, in CRC through the TCGA database. We found that HNF4α had the highest correlation with TCF1 and TCF3, but not LEF1 and TCF4 (Figure 4A-4D). Meanwhile, we individually knocked down these transcription factors in SW480 (Figure 4E-F). Western bot analysis showed that HNF4α was reduced in TCF3 knockdown cells but not in shRNA-TCF1, shRNA-TCF4 or shRNA-LEF1 treated cells (Figure 4E). We further confirmed the result in two different CRC cell lines, LS174T and DLD1. Similarly, TCF3 knockdown decreased HNF4α expression (Figure 4G). Moreover, ectopic expression of TCF3 induced HNF4α level, but ectopic expression of TCF3 in β-catenin shRNA cells can’t rescue HNF4α level (Figure 4H-I). These results suggest that TCF3 is a major transcriptional factor that mediates the action of Wnt/β-catenin in induction of HNF4α in CRC.

**Figure 4.**
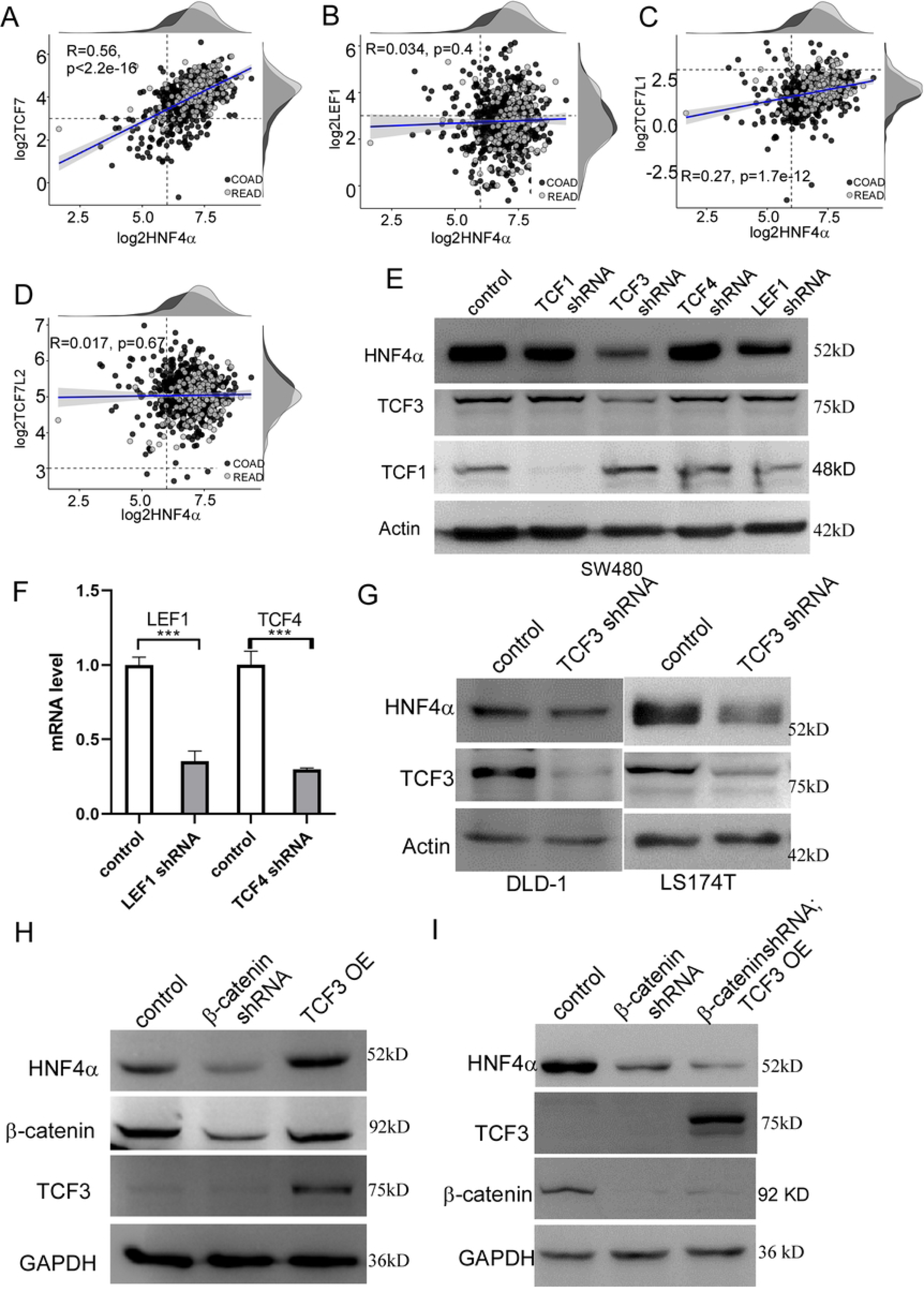
TCF-3 is a major transcription factor to induce HNF4α expression. A-D. Correlation analysis of mRNA levels between HNF4α & TCF/LEF family members using TCGA database. E. Western blotting analysis of the effects of TCF/LEF members on HNF4α expression. F. Q-PCR analysis of the shRNA efficiency on LEF1 and TCF4 G. Western blot analysis of the effect of TCF3 knockdown on HNF4α levels in DLD1 and LS174T cell lines. H. Western blotting analysis of the effect of β-catenin knockdown and TCF3 overexpression on HNF4α expression. I. Western blotting analysis of the effect of TCF3 overexpression in β-catenin knockdown cells on HNF4α expression.

### 5. TCF3 directly binds to the -90--80bp region of the HNF4α promoter

To investigate whether TCF3 can directly bind to the HNF4α promoter to regulate HNF4α transcription, we first examined the TCF3 consensus binding motif “CACCTGC” in the HNF4α promoter region through the website (https://jaspar.genereg.net/). As shown in Figure 5A, there are 10 TCF3 putative binding sites within a 2.0-kb HNF4α promoter. To determine which binding site(s) is required for TCF3-mediated HNF4α transcription, we constructed deletion mutants of the HNF4α promoter including pGL3-P_HNF4α_1700bp, pGL3-P_HNF4α_800bp and pGL3-P_HNF4α_300bp, which contain 10, 7 and 3 TCF3 putative binding sites, respectively (Figure 5A). Luciferase reporter assays revealed that knockdown of TCF3 significantly reduced the basal promoter activity in three HNF4α deletion promoter mutants (Figure 5B), suggesting that pGL3-P_HNF4α_300bp contains major TCF3 response motifs (Figure 5A). Since there are 3 TCF3 putative binding sites in pGL3-P_HNF4α_300bp, we next mutated each site in this promoter region (Figure5C). Interestingly, mutations of the 90 to 80-bp motif, which is the closest to transcription start site of HNF4α gene, completely abrogated the pGL3-P_HNF4α_300bp promoter activity, whereas mutation of the rest two sites had no significant effect on the promoter activity (Figure 5D). These findings suggest that TCF3 transcriptionally activates HNF4α primarily through binding to the first motif of the HNF4α promoter.

**Figure 5.**
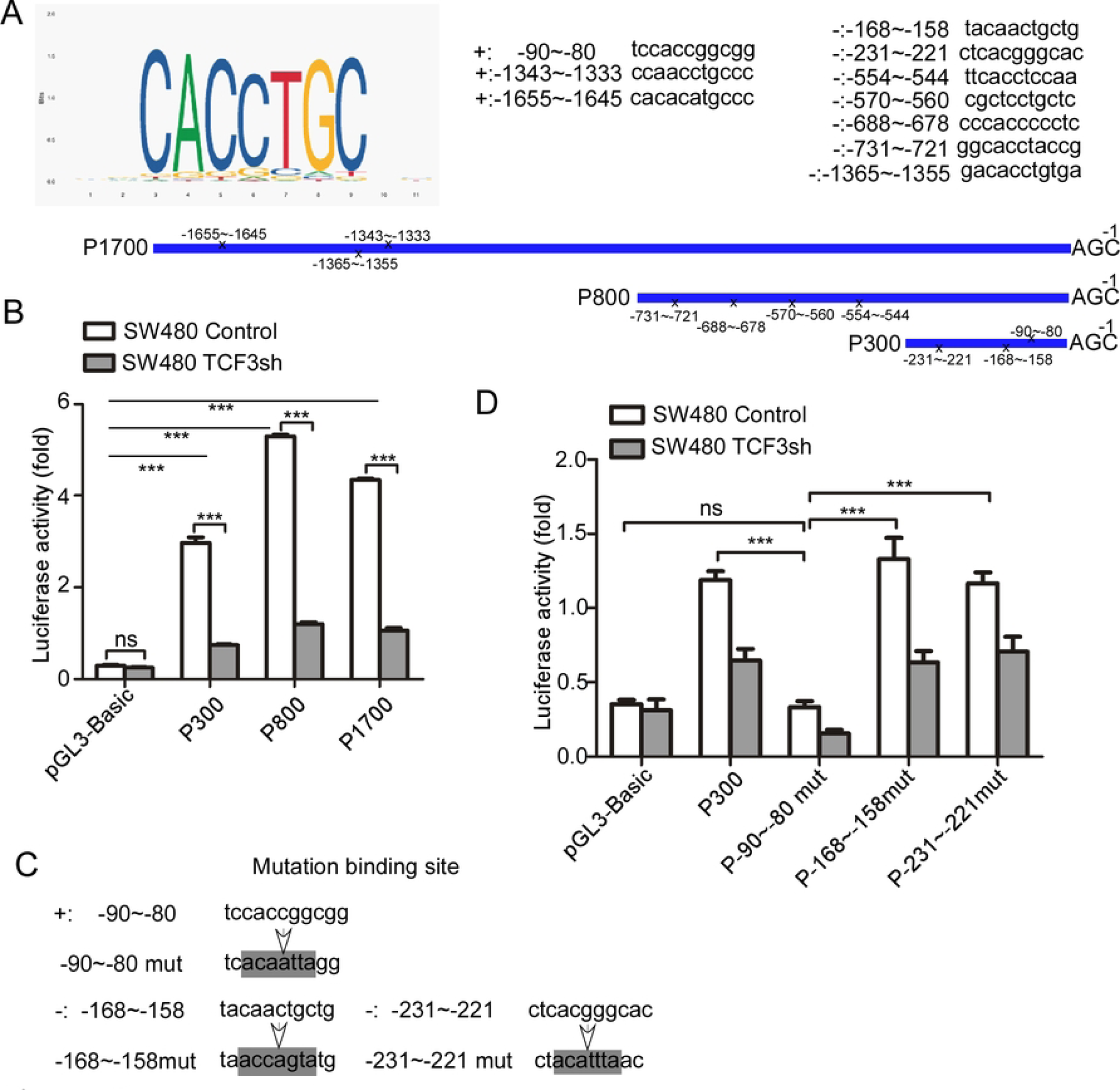
TCF3 directly binds to the HNF4α promoter and regulates its transcription. A. JASPAR (https://jaspar.genereg.net/) analysis of 1.7-kb HNF4α promoter for putative TCF3 transcription factor binding motif(s). Graphical representation of predicted binding sites and TCF3 binding sequence motifs shown on the top left. B. Luciferase activity of HNF4α promoters. C. Mutation of each TCF3 binding motif in the pGL3-HNF4α/300bp promoter region. D. Luciferase activity of three mutants of pGL3-HNF4α/300bp promoter. ***p<0.001.

### 6. Overexpression of HNF4α activates WNT and promotes β-catenin nuclear localization

We next performed RNA sequencing analysis between SW480-vector and SW480-HNF4α cells in an attempt to investigate the mechanism of HNF4α in colorectal tumorigenesis. The signals enriched by HNF4Α overexpression were involved in cardiac system diseases, neurological diseases, tumor progression and other features, among which the most tumor-related signaling pathways were the MAPK, Hippo and Wnt signaling cascades (Figure 6A). Notably, Wnt1, Wnt4, Wnt7b and Wnt11 were up-regulated in SW480-HNF4α cells when compared to SW480-vector control cells (Figure 6B), which were further confirmed by qPCR assay (Figure 6C). However, we did not observe expression level changes of β-catenin in the cells ectopically expressing HNF4α (Figure 6D). As HNF4α induces expression levels of several Wnt family members, we hypothesized that HNF4α could promote β-catenin translocation from the cytoplasm to the nucleus. Cell fractionations were obtained from SW480-vector and SW480-HNF4α cells. Immunoblotting analysis revealed that the nuclear fraction of β-catenin was significantly enriched in the HNF4α overexpressing cells (Figure 6E). Collectively, our findings indicate that HNF4α and Wnt/β-catenin regulated each other and form a feedback regulation loop to contribute to colorectal carcinogenesis.

**Fig 6.**
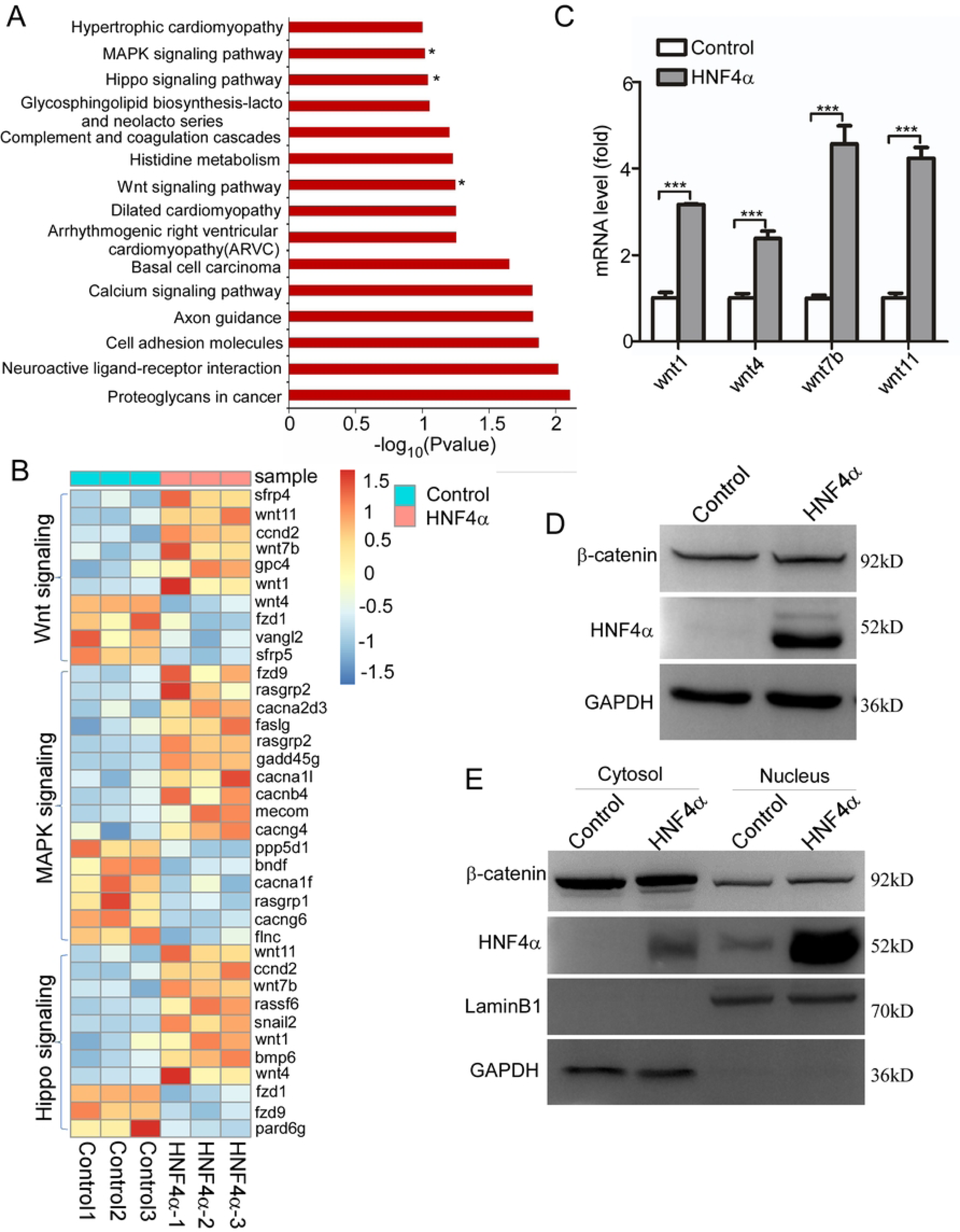
HNF4α induces expression of Wnt family members and β-catenin translocation from the cytoplasm to the nucleus. A. KEGG pathway analysis by RNA sequencing data obtained from SW480-vector control and SW480-HNF4α cells. B. A heat map showing expression levels of the genes involved in WNT, MAPK, Hippo signaling pathways that are regulated by HNF4α. C. q-PCR analysis of HNF4α overexpression on Wnt family members level. D. Western blot analysis of the effect of HNF4α overexpression on β-catenin level E. Western blot analysis of the effect of HNF4α overexpression on β-catenin nuclear localization.

### 7. The β-catenin/TCF3-HNF4**α** axis correlates with colorectal tumorigenesis

To evaluate the clinical significance of β-catenin/TCF3-HNF4α in colorectal cancer, we performed IHC analysis and examined the HNF4α and β-catenin/TCF3 expression level in a colon cancer tissue array and investigated the correlation of β-catenin/ TCF3 with HNF4α in colorectal cancer. The results showed that the tumor tissue with high levels of β-catenin/ TCF3 usually express high levels of HNF4α, and the tumor tissue with low levels of β-catenin/ TCF3 usually have low levels of HNF4α (Figure 7A). The correlation analysis indicated that HNF4α was significantly correlated with β-catenin and TCF3 (Figure 7B and 7C). Taken together, these results indicated that HNF4 was correlated with the WNT/β-catenin pathway and involved in colorectal tumorigenesis.

**Fig7.**
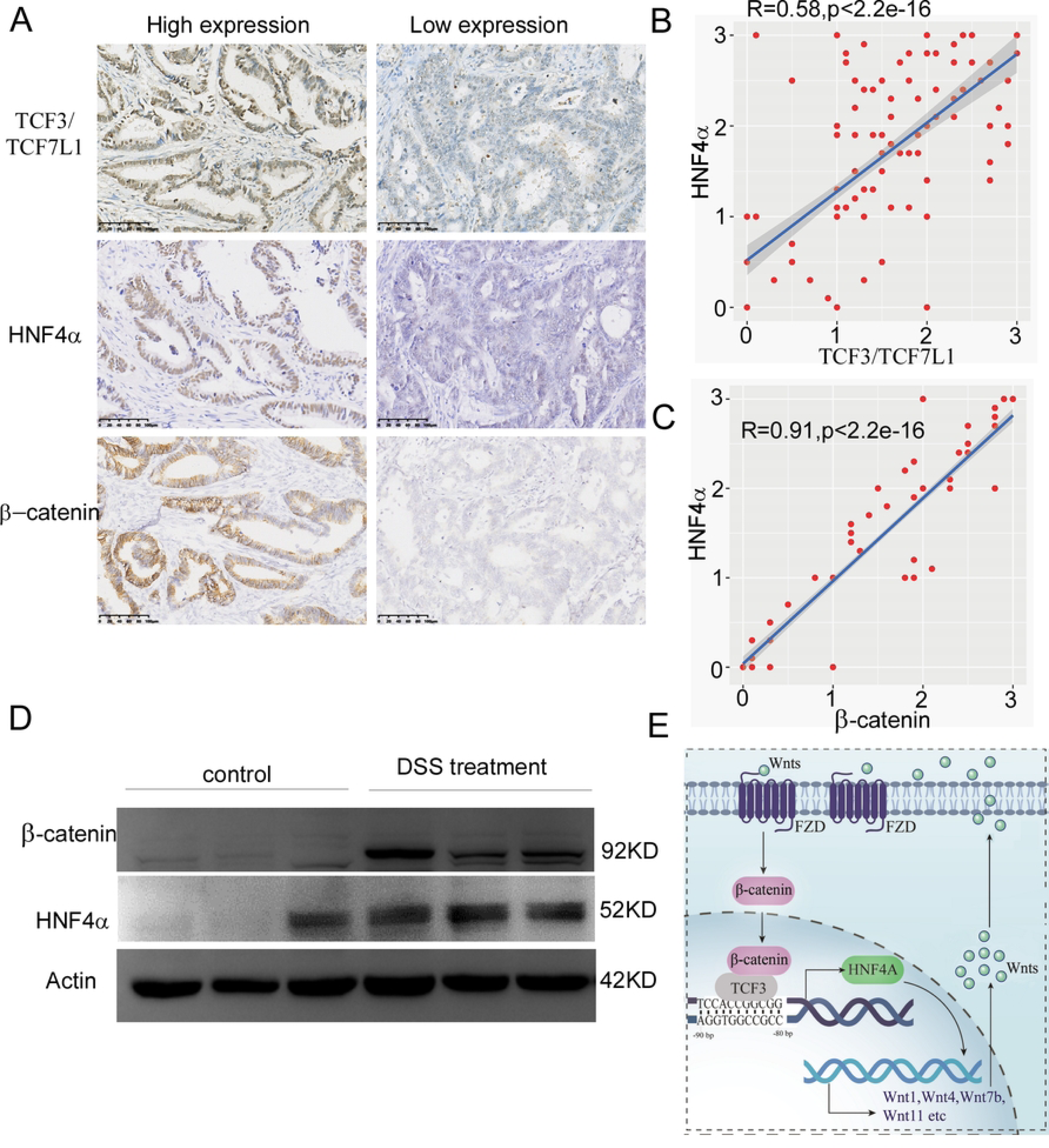
β-catenin/TCF3-HNF4α axis correlate with colorectal tumorigenesis. A. Representative images of β-catenin, TCF3 and HNF4α immunohistochemical staining in colorectal cancer tissue B. Correlation analysis between TCF3 and HNF4 α level C. Correlation analysis between β-catenin and HNF4 α level D. Western blot analysis of β-catenin, TCF3 and HNF4α level in colon tissue of AOM-DSS mouse colon cancer model E. Model of β-catenin/TCF3-HNF4α regulation in colorectal tumorigenesis

To further test our hypothesis, we employed AOM/DSS Model of Colitis-Associated colorectal Cancer model[21], and AOM/DSS treatment can not only induce intestinal inflammation and colorectal cancer, but also induce highly express β-Catenin and HNF4 in the intestinal tissues of treated mice (Figure 7D). The result implied that the β-catenin/ TCF3-HNF4α axis plays an important role in colorectal cancer (Figure 7E).

In conclusion, our results show that HNF4α acts as a WNT/β-catenin downstream factor and affected by direct regulation of β-catenin/TCF3 to regulate the occurrence of colorectal cancer. Overexpression of HNF4α can activate the expression of wnt family members and promote β-catenin nuclear localization and promotes wnt/β-catenin signal pathway. The Wnt/β-catenin-TCF3-HNF4α feedback loop promotes colorectal tumorigenesis. This study can provide strategies for the treatment and prevention of colorectal cancer.

## Discussion

CRC is a highly heterogeneous disease, the roles of HNF4α in colorectal tumorigenesis have been reported are contradictory, Single-cell chromatin accessibility analysis reveals that HNF4α is key iCMS-specific transcription factors in colorectal malignant tumor cells, suggesting that HNF4α may play important roles in colorectal tumorigenesis and progression[22]. However, the role and regulatory mechanism of HNF4α in colorectal cancer are still largely unknown.

Wnt/β-catenin activation is a recognized driver of colorectal carcinogenesis, however, the regulation of wnt/ β-catenin in colorectal cancer is still largely unknown. HNF4α is necessary for liver development and function and is a negative regulator for liver cancer. Previous research reports have shown that HNF4α expression was negatively regulated by Wnt-β-catenin signaling in hepatocellular carcinoma[23]. The regulatory relationship between Wnt/β-catenin and HNF4α in colorectal cancer is rarely reported.

Here we showed that HNF4αpromotes the development of colon cancer. HNF4α is a Wnt/β-catenin effector and is directly regulated by Wnt/β-catenin signaling. HNF4α overexpression further affects Wnt/β-catenin signaling pathway activation and promotes the colorectal cancer. There are four transcription factors, TCF1, LEF1, TCF3 and TCF4) in the wnt/β-catenin signaling pathway and they exhibit significant differences in regulating target genes[24]. Compared to the other three transcription factors, TCF-3 was the least reported in tumors. The mutual regulatory relationship between TCF3 and HNF4α has not been reported. Our study indicates that β-catenin activates TCF3, which directly binds to the HNF4α promoter region and regulates HNF4α expression and colorectal tumorigenesis.

In conclusion, our study demonstrated that the Wnt/β-catenin-TCF3-HNF4α feedback loop confers colorectal tumorigenesis. Our result also suggested that HNF4α is a new target of Wnt/β-catenin-TCF3(TCF7L1) and HNF4α overexpression feedback regulates WNT expression and promotes β-catenin nuclear localization. Further investigation of the role and the specific mechanism of HNF4α in the development and progression of CRC is of great significance for establishing HNF4α as a therapeutic target in CRC. Our study will provide strategies for the treatment and prevention of colorectal cancer.

## Materials and Methods

### Cell culture and activator treatment

Colon cancer cell line SW480, SW620 and DLD-1 were cultured in RPMI1640 medium supplemented with 10% fetal bovine serum (FBS) and 1% penicillin/streptomycin. For the inhibitor and activator treatment, cells (5 × 10^5^) were seeded in 6 well plates and treated with 2uM or 5Um SKL2001 and CHIR99021 for 12 h, 24 h, and 36 h after the cells attached to the dish.

### RNA interference

RNA interference was performed using the plko.1 vector. To efficiently knockdown β-catenin and TCF family, the shRNAs targeting to different genes were synthetized and inserted to plko.1 vector between Age I and EcoR I. The target sequences were as follows:

TTGTTATCAGAGGACTAAATA (β-catenin shRNA) ;

GCACCTACCTGCAGATGAAAT (TCF3 shRNA);

CAACTCTCTCTCTACGAACAT (TCF1 shRNA);

CCATCAGATGTCAACTCCAAA (LEF1 shRNA);

CCTTTCACTTCCTCCGATTAC (TCF4 shRNA);

### Western blotting

Cells were lysed in RIPA buffer (20 mM Tris-HCl pH 7.5, 100 mM NaCl, 0.1% SDS, 0.5% sodium deoxycholate, and 1 mM PMSF) containing Complete Protease Inhibitor Cocktail (Roche). Cell lysates were spun down at 12,000 rpm for 10 min, and 50 µg supernatants were resolved on SDS-PAGE and blotted with the indicated antibodies. The amount of GAPDH was used as the loading control. The results were detected by an ECL-plus Western blotting detection system (Tanon-5200Multi). The primary antibodies used in this study were as follows: HNF4α, Rabbit #3113 CST WB: 1:1000 TCF3, Rabbit #PA5-40327 Thermo Fisher Tech. WB: 1:1000 TCF1, MOUSE #sc-271453 Santa cruz bio. WB: 1:1000 β-catenin, Rabbit # A117811 GeneScript WB: 1:1000 Actin, Rabbit #4970 CST WB: 1:1000 GAPDH, mouse#sc-166574 Santa cruz bio. WB: 1:1000

### Quantitative reverse transcription–polymerase chain reaction (qPCR)

Total RNA was extracted using Trizol (Invitrogen) and chloroform. 2 μg of RNA was used astemplate to generate cDNAs using the ImProm-II Reverse Transcription system (Promega, Madison, Wisconsin, USA). qPCR reactions were carried out on an MX3000P system (AgilentTechnologies, Santa Clara, CA).

### Mouse Xenograft

All the mice were obtained from the Animal Research and Resource Center, Yunnan University. {Certification NO. SCXK(Dian)K2021-0001}. All animal experiments were performed according to protocols approved by the Animal Care Committee of the Yunnan University (Kunming, China).

For the Subcutaneous tumor models,1×10^6^ SW620 cells were re-suspended in 100 μl 0.25 mg/ml Matrigel (Corning) with PBS buffer, then injected into the flanks of the nude mouse. The tumor volumes were measured from day 3 to 11 after injection. At 11 days after the injection, tumors were dissected. Tumors were then imaged and weighted.

### GESA analysis and correlation analysis

The mRNA sequencing data of COAD READ BRCA LIHC LUAD LUSC PAAD SKCM were download from TCGA database (https://portal.gdc.cancer.gov/repository) respectively, and the merge normal tissues UCSC Xena (https://xenabrowser.net) mRNA Sequencing data was downloaded as control. the Pearson correlation coefficient R value and the correlation curve were calculated and draw by Using the ggplot2 function in R 4.2.0 software. the boxplot was drawn using the simplevis package after processing the TPM as log2 (TPM+1). The genes that significantly(p<0.01, r ≥ 0.3 or r ≤ - 0.3) related to HNF4 α were screened by FPKM value of COAD READ and used for GSEA and KEGG enrichment analysis[25].

### Luciferase Assay

the Dual Luciferase Assay were performed according to a previously reported protocol with minor modification [2013 Jianwei sun JBC]. The HNF4 α promoter luciferase reporter constructs were generated by inserting 1.7K, 800bp or 300bp human HNF4α promoter into pGL3 basic vector (Promega) between XhoI and HindIII. To perform dual luciferase reporter assay, 2,0000 SW480 cells were seeded in 12-well plates and cultured overnight. Cells were transfected with 1 g/well HNF4 α promoter reporter together with 100 ng/well Renilla luciferase construct (pRL-TK) using Lipofectamine 2000. 24h after transfection, the cells were was lysed and Cell lysates were subjected to dual reporter luciferase assays according to the manufacturer’s instructions (Promega).

### Tissue microarray and immunohistochemistry

Tissue microarrays (TMAs) were constructed using a manual tissue microarray instrument (Beecher Instruments) equipped with a 2.0 mm punch needle, as we previously described. An immunohistochemical (IHC) study of rabbit anti-human β-catenin, rabbit anti-human TCF3/TCF7L1 antibody, rabbit anti-human HNF4α antibody was carried out on formalin-fixed paraffin, with a 4-μm-thick serial section of tissue, according to the manufacturer’s recommended protocol. The levels of β-catenin, TCF3/TCF7L1 and HNF4α were assessed via the average of 5 count fields per patient in the original magnification of X200 on light microscopy. The positive cells for HNF4α were defined as those with brown staining. The expression of β-catenin, TCF3/TCF7L1 and HNF4α was scored based on staining intensity. Staining intensity was subclassified as follows: 0, negative; 1, weak; 2, moderate; and 3, strong.

The primary antibodies used in this study were as follows:

HNF4α, Rabbit #3113 CST IHC: 1:200

TCF3, Rabbit #PA5-40327 Thermo Fisher Tech. IHC: 5 µg/mL

β-catenin, Rabbit # ZA-0646 Zhongshan Golden Bridge Biotech. IHC: ready to use

### Statistics analysis

Data were analyzed with Prism (GraphPad software). Statistical analyses were performed using *t*-tests or ANOVA. * P*<*0.05 was considered statistically significant. ** P*<*0.01 was considered significant. *** P*<*0.001 was considered extremely significant. P*>*0.05 was considered not significant (ns)

## Acknowledgments

We thank Dr. Jing Li for their helpful suggestion and comments on the manuscript. This work was supported by the National Natural Science Foundation of China (NSFC) fund (82273460 and 32260167), the Yunnan Applied Basic Research Projects (202101AV070002 and 202301AS070117), and grants (Grant No. 2023Y0222, KC-

23234451, KC-23233927 and 202310673059) from Yunnan University.

## Authors’ Contributions

LS, WB, CL and RD: Experiments and data analysis. QH, YZ, XY, and RS: Vector construction and dual-luciferase assay. XL, MM and JY: Cell culture and Western blot. JS, YS and JS: Experiments design, data analysis, manuscript writing. All authors read and approved the final manuscript.

